# The eukaryotic Initiation Factor 6 (eIF6) regulates ecdysone biosynthesis by modulating translation in *Drosophila*

**DOI:** 10.1101/201558

**Authors:** Arianna Russo, Guido Gatti, Roberta Alfieri, Elisa Pesce, Kelly Soanes, Sara Ricciardi, Marilena Mancino, Cristina Cheroni, Thomas Vaccari, Stefano Biffo, Piera Calamita

**Author notes:** These authors contributed equally to this work (Stefano Biffo, Piera Calamita). Address correspondence to Piera Calamita or Stefano Biffo.

## Abstract

During development, ribosome biogenesis and translation reach peak activities, due to impetuous cell proliferation. Current models predict that protein synthesis elevation is controlled by transcription factors and signalling pathways. Developmental models addressing translation factors overexpression effects are lacking. Eukaryotic Initiation Factor (eIF6) is necessary for ribosome biogenesis and efficient translation. *eIF6* is a single gene, conserved from yeasts to mammals, suggesting a tight regulation need. We generated a *Drosophila melanogaster in vivo* model of eIF6 upregulation, demonstrating a boost in general translation and the shut off of the ecdysone biosynthetic pathway. Translation modulation in S2 cells showed that translational rate and ecdysone biosynthesis are inversely correlated. In vivo, eIF6-driven alterations delayed programmed cell death (PCD), resulting in aberrant phenotypes, partially rescued by ecdysone administration. Our data show that eIF6 triggers a translation program with far-reaching effects on metabolism and development, stressing the driving and central role of translation.

## INTRODUCTION

During cell proliferation, ribosomal proteins (RPs) and eukaryotic Initiation Factors (eIFs) are necessary and in high demand for ribosome biogenesis and translation ^1-5^. Proteins involved in ribosome biogenesis do not usually have a role in the translational control and *vice versa* ^6^. However, the eukaryotic Initiation Factor 6 (eIF6) is remarkably unique ^7^: a nuclear pool is essential for nucleolar maturation of the 60S large subunit ^8^, while cytoplasmic eIF6 acts as a translation factor. Mechanistically, eIF6 is an anti-association factor: by binding to the 60S subunit, eIF6 prevents its premature joining with a 40S not loaded with the preinititiaton complex. Release of eIF6 is then mandatory for the formation of an active 80S ^9^. In mammals, eIF6 translation activity increases fatty acid synthesis and glycolysis through translation of transcription factors such as CEBP/β, ATF4 and CEBP/δ containing G/C rich or uORF sequences in their 5’UTR ^10,11^. The dual action of eIF6 in ribosome biogenesis and translation suggests that it may act as a master gene regulating ribosomal efficiency. Remarkably, point mutations of eIF6 can revert the lethal phenotype of ribosome biogenesis factors such as SBDS^12^ and eFL1p^13^. eIF6 is highly conserved in yeast, *Drosophila* and humans ^14^. During evolution, the *eIF6* gene has not been subjected to gene duplication. Despite its ubiquitous role, eIF6 levels *in vivo* are tightly regulated, showing considerable variability of expression among different tissues ^15^. Importantly, high levels of eIF6 or hyperphosphorylated eIF6 are observed in some cancers ^16,17^. eIF6 is rate-limiting for tumor onset and progression in mice ^18^. In addition, eIF6 amplification is observed in luminal breast cancer patients ^19^ and may affect also cancer cell migration ^20,21^. However, whether eIF6 overexpression *per se* can change a transcriptional program in the absence of other genetic lesions is unknown.

To determine the effects of *eIF6* increased gene dosage *in vivo*, we took advantage of *Drosophila melanogaster*, an ideal model to manipulate gene expression in a time and tissue-dependent manner, using the GAL4/UAS system ^22,23^. We reasoned that a gain of function approach could allow us to evaluate the effects of eIF6 overexpression in the context of an intact organ. To this end, we used the fly eye, an organ not essential for viability, whose development from epithelial primordia, the larval eye imaginal disc, is well understood. The adult fly compound eye is a stunningly beautiful structure of approximately 800 identical units, called ommatidia ^24^. Each ommatidium is composed of eight neuronal photoreceptors, four glial-like cone cells and pigment cells ^25,26^. By increasing eIF6 levels in the eye, we have found alterations in physiological apoptosis at the pupal stage, correlating with an increase in general translation. Importantly, we also observed a reshaping of the eye transcriptome that revealed a coordinated downregulation of the ecdysone biosynthesis pathway. Overall, our study provides the first *in vivo* evidence that an increase in translation, dependent on a heightened *eIF6* gene dosage, may drive metabolic changes and a transcriptional rewiring of a developing organ. Our model shows that overexpression of a translation factor *per se* induces a gene expression program and stresses the central role of translational control.

## RESULTS

### Increased eIF6 levels cause embryonic lethality and aberrant morphology

Regulation of eIF6 levels is stringent in normal conditions^15^ with evidence for eIF6 amplification^19^ and overexpression in cancer^14,16,27-29^. We used the *Drosophila melanogaster* model to establish whether an increased dosage of eIF6 could drive specific developmental decisions.

We first assessed the effects caused by the loss of the *Drosophila* homologue of *eIF6*. To this end, we used the *P* element allele *eIF6^k13214^* ^30^, inducing mitotic clones homozygous for *eIF6^k13214^* in first instar larvae by heat shock-induced FLIP/*FLP-*mediated homologous recombination ^31^. We did not observe clones of eIF6 mutant cells with the exception of small ones in the wing margin. Similar results were obtained in a *minute (M)* background that provides a growth advantage to mutant cells, or by targeted expression of FLP in the wing margin (Supplementary Fig. 1a). Together, these results indicate that eIF6 is required for cell viability in *Drosophila*, as previously observed in yeast ^17^ and mammals ^8^, precluding significant studies on the effects of eIF6 inhibition, a phenomenon anyhow absent in physiological conditions.

Next, we assessed the effects of eIF6 gain of function, which is often observed in several cancers, by expressing *eIF6* ubiquitously using the *TubGAL4* driver. Ectopic expression resulted in late embryonic lethality (Supplementary Fig. 1b), suggesting that increased levels of eIF6 dramatically disrupt gene expression. To circumvent early lethality, we then focused on a non-essential fly organ, the eye. Increased eIF6 expression during late larval eye disc development, driven by the *GMRGAL4* driver (*GMR>eIF6*), causes the formation of a reduced and rough adult eye (Fig. 1a). Using a new antibody specific for *Drosophila* eIF6 that we developed (see Material and Methods section) we estimated that the level of expression was about doubled compared to controls (Fig. 1b). SEM analysis showed severe disruption of the stereotypic structure of the wild-type eye, with flattened ommatidia and bristles arranged in random patterns (Fig. 1c). Semithin sections evidenced that the phenotype correlates with an aberrant arrangement and morphology of the eye cells (Fig. 1d). These data show that doubling the *eIF6* gene dosage in the *Drosophila* eye causes disruption of eye development.

**Figure 1.**
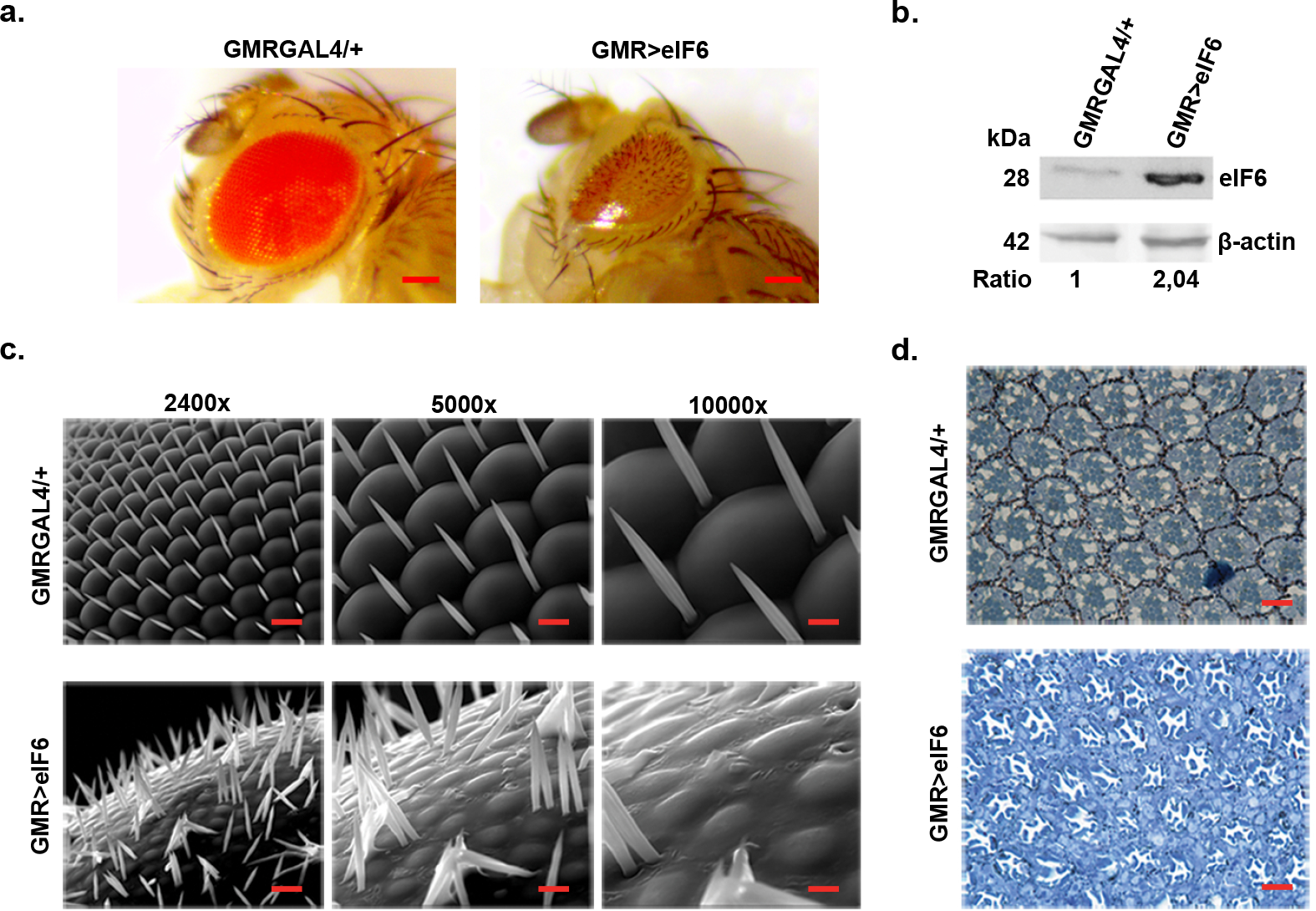
Increased eIF6 levels in the developing eye result in a rough eye phenotype. **(a)** Representative stereomicroscope images of *GMRGAL4/+* and *GMR>eIF6* eyes, showing a noteworthy rough eye phenotype. Scale bar 300 µm. **(b)** Western blot showing the levels of eIF6 expression in *GMRGAL4/+* and *GMR>eIF6* adult eyes. Representative western blots from three independent experiments are shown. Molecular weight markers (kDa) are shown to the left of each panel. Ratio was calculated with ImageJ software. The value corresponds to the intensity ratio between eIF6 and β-actin bands for each genotype. **(c)** Representative SEM images of *GMRGAL4/+* and *GMR>eIF6* adult eyes. eIF6 overexpressing eyes have an aberrant morphology, showing flattened ommatidia and randomly arranged bristles. Scale bar, respectively for 2400X, 5000X and 10000X magnifications are 10 µm, 5 µm and 2.5 µm **(d)** Representative tangential sections of *GMRGAL4/+* and *GMR>eIF6* adult eyes indicating that photoreceptors are still present in *GMR>eIF6* eyes, even if their arrangement is lost. Scale bar 10 µm.

### Increased *eIF6* gene dosage delays physiological apoptosis

To understand the origin of the defects observed in *GMR>eIF6* adult eyes, we analyzed eye development in larvae, starting from the third instar, the stage at which the GMR-GAL4 driver starts to be expressed. We found that third instar imaginal discs with higher levels of eIF6 showed no differences in terms of morphology or cell identity, when compared to controls (Supplementary Fig. 2a). Then, we analyzed pupal development. In *GMR>eIF6* flies at 40h after puparium formation (APF) both neuronal and cone cells were present in the correct numbers. However, ommatidial morphology was altered (Supplementary Fig. 2b). We considered that a fundamental event controlling ommatidial morphology is the developmentally-controlled wave of Programmed Cell Death (PCD), sweeping the tissue from 25h to 42h APF ^26^. Thus, we analyzed by immunostaining the expression of *Drosophila* apoptotic effector caspase Dcp-1, as a marker of PCD, at 40h APF. Control retinae showed clear presence of apoptotic cells. Remarkably, apoptotic cells were completely absent in *GMR>eIF6* retinae (Fig. 2a).

**Figure 2.**
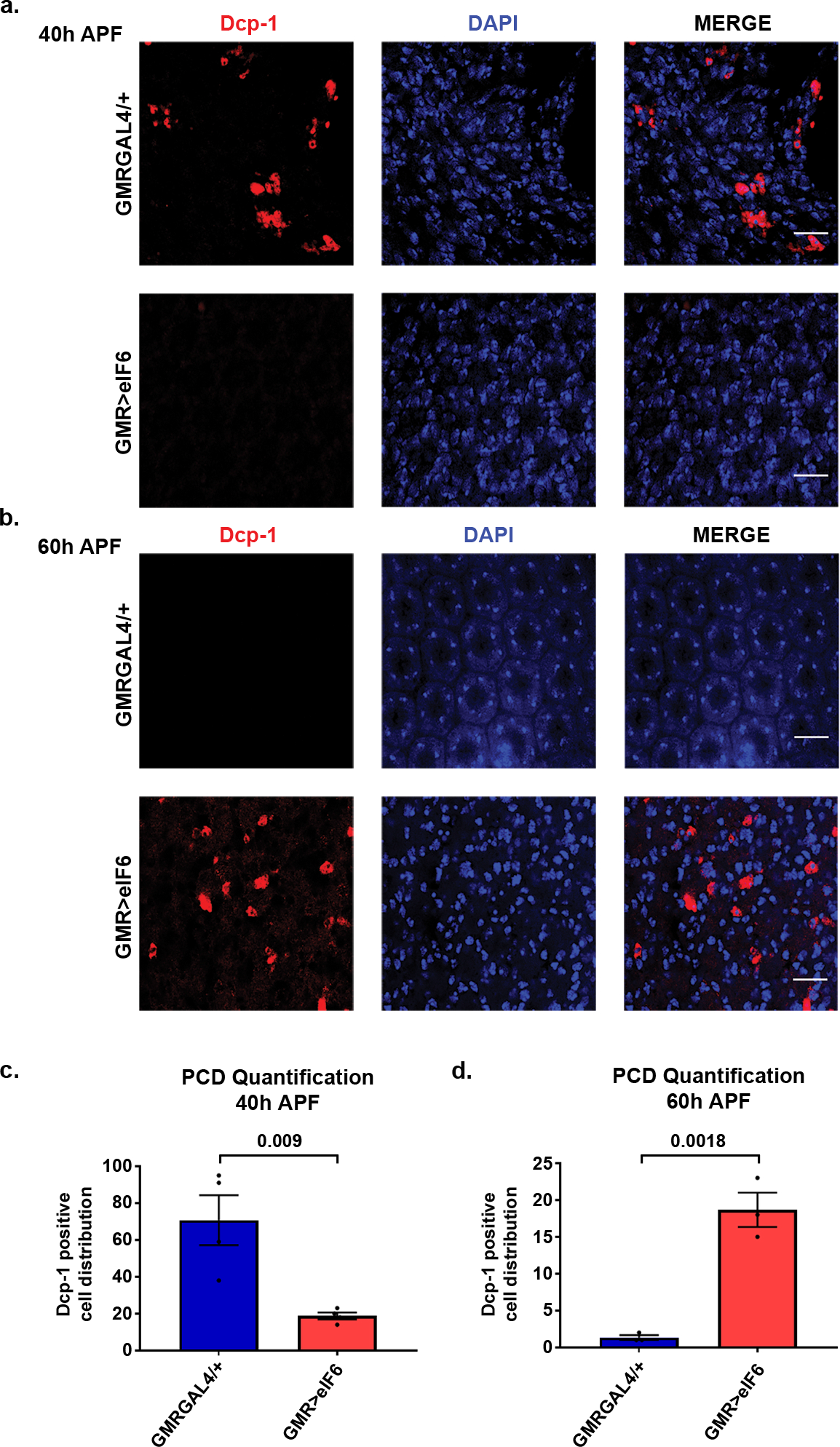
The apoptotic wave is delayed when *eIF6* gene dosage is increased. **(a)** Mid-pupal stage retinae (40h APF) stained for the *Drosophila* caspase Dcp-1. *GMRGAL4/+* retinae show Dcp-1 positive cells, indicating that PCD is ongoing at this developmental stage. On the contrary, *GMR>eIF6* retinae do not show Dcp-1 positive cells, indicating a block in PCD. Scale bar 10 µm. **(b)** Late-pupal stage (60h APF) retinae stained for the *Drosophila* caspase Dcp-1. *GMRGAL4/+* retinae show the absence of Dcp-1 positive cells, as expected (PCD already finished at this developmental stage). On the contrary, *GMR>eIF6* retinae, show Dcp-1 positive cells, indicating a delay in PCD associated to more eIF6 levels. Scale bar 10 µm. **(c-d)** Representative barplot showing the Dcp-1 positive cells counts average from four different area (n=4) at 40h APF **(c)** and 60h APF **(d)** retinae with error bars indicating the SEM. P-values were calculated using an unpaired two-tailed Student t-test. Dcp-1 positive cells counts indicate an overall delay and increase in PCD when *eIF6* gene dosage is increased during eye development.

Dcp-1 positive cells, i.e. apoptotic cells, appeared in *GMR>eIF6* retinae only at 60h APF (Fig. 2b). In contrast, 60h APF wild-type retinae did not show any longer apoptotic cells (Fig. 2b). Quantitation of the number of Dcp-1 positive cells at 40h APF and 60h APF in *GMR>eIF6* (Fig 2c) revealed up to 75% reduction in the number of apoptotic cells at 40h APF. A change in apoptosis dynamics was also visualized by TUNEL assay at 28h APF, the time at which PCD starts in control retinae. Here, we observed the absence of apoptotic nuclei in the *GMR>eIF6* retinae, while *GMRGAL4/+* retinae showed several (Supplementary Fig. 2c). We stained for the *Drosophila* β-catenin homologue Armadillo (Fig. 3), which localizes to membranes of cells surrounding photoreceptors, providing an indication of their number. At 40h APF, control retinae presented the typical staining expected for Armadillo, while *GMR>eIF6* retinae showed the presence of extra-numerary cells around the ommatidial core (Fig. 3a). This finding is in line with the possibility that interommatidial cells (IOCs) were not removed by PCD. By counting the number of cells in each ommatidium, we determined that *GMR>eIF6* retinae possess more than 15 cells, corresponding to approximately 30% more than that of a wild-type ommatidium (Supplementary Fig. 3a). Later in development, both at 60h and at 72h APF, in *GMR>eIF6* retinae Armadillo was no longer detectable, while in wild-type retinae the pattern of Armadillo was maintained (Fig. 3b and Supplementary Fig. 3b). These data indicate that late PCD in *GMR>eIF6* is likely to inappropriately remove most inter-ommatidial cells (IOCs). In conclusion, the first effect of eIF6 high levels is not a cytotoxic effect, but a block of apoptosis that leads in turn to a disrupted developmental program.

**Figure 3.**
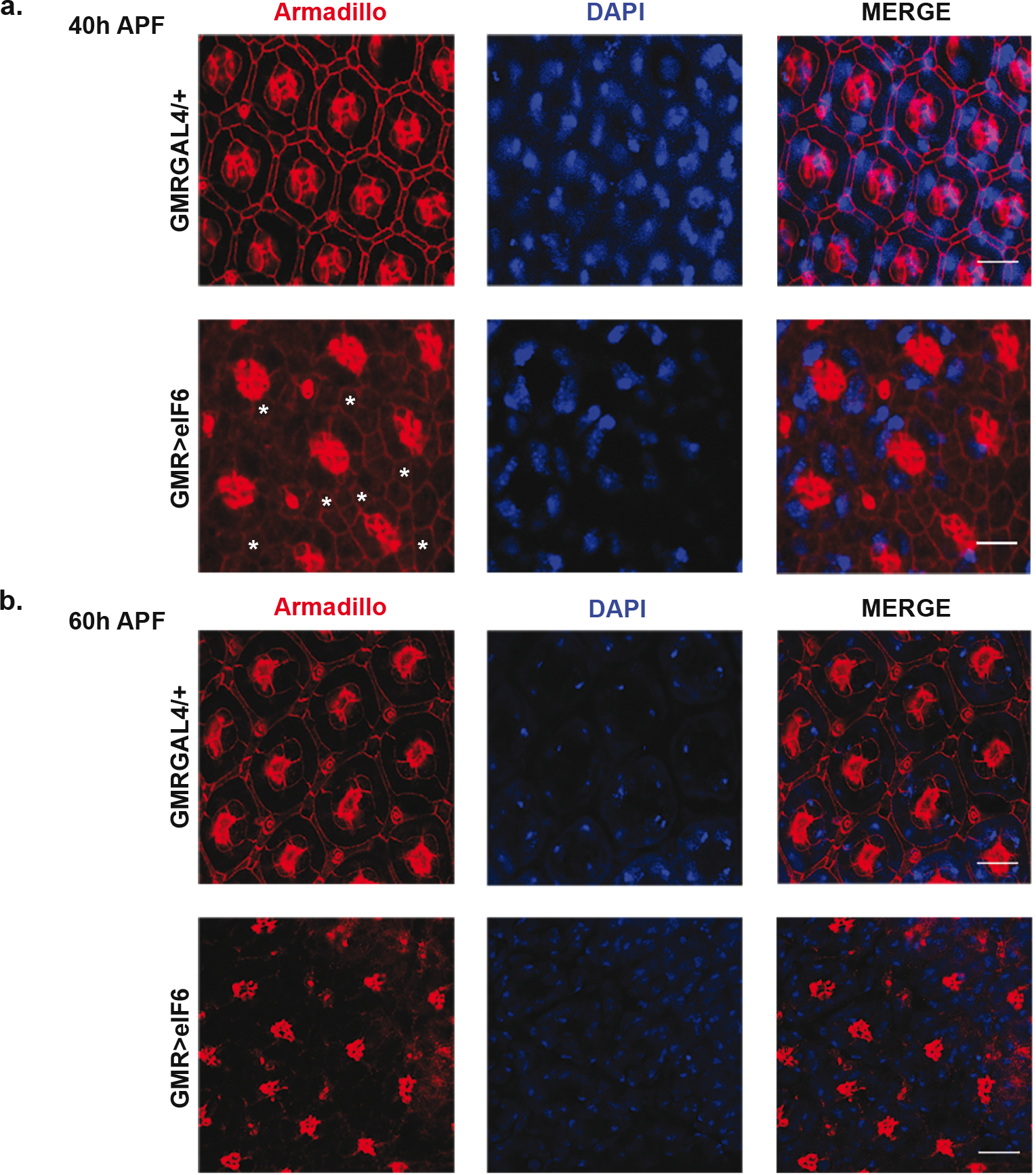
Cell number is altered during pupal stage in *GMR>eIF6* retinae. **(a)** Mid-pupal stage (40h APF) retinae stained for Armadillo, the *Drosophila* β-catenin homologue, showing that when eIF6 is increased there are extra-numerary cells (indicated as *) around each ommatidium. Scale bar 10 µm. **(b)** Late-pupal stage (60h APF) retinae stained for Armadillo, showing the loss of all cells around ommatidia upon eIF6 overexpression. Scale bar 10 µm.

### Increased *eIF6* dosage in cone cells is sufficient to delay apoptosis

Cone cells and IOCs are known to be the main actors during physiological PCD^32^. Thus, to understand whether increased eIF6 levels restricted to cone cells affects eye morphology we overexpressed eIF6 under the control of the cone cell specific driver, *spaGAL4*. We observed a similar phenotype to that of *GMR>eIF6*, albeit a milder one (Fig. 4a-b and Supplementary Fig. 4a). Importantly, eIF6 overexpression specifically in cone cells, caused absence of Dcp-1 staining in 40h APF retinae (Fig. 4c and Supplementary Fig. 4b), confirming a block in apoptosis. In contrast, apoptosis was evident at 60h APF (Fig. 4d), in line with what we observed in *GMR>eIF6* retinae. Thus, the expression of eIF6 in cone cells is sufficient to alter PCD and cause defects in eye development.

**Figure 4.**
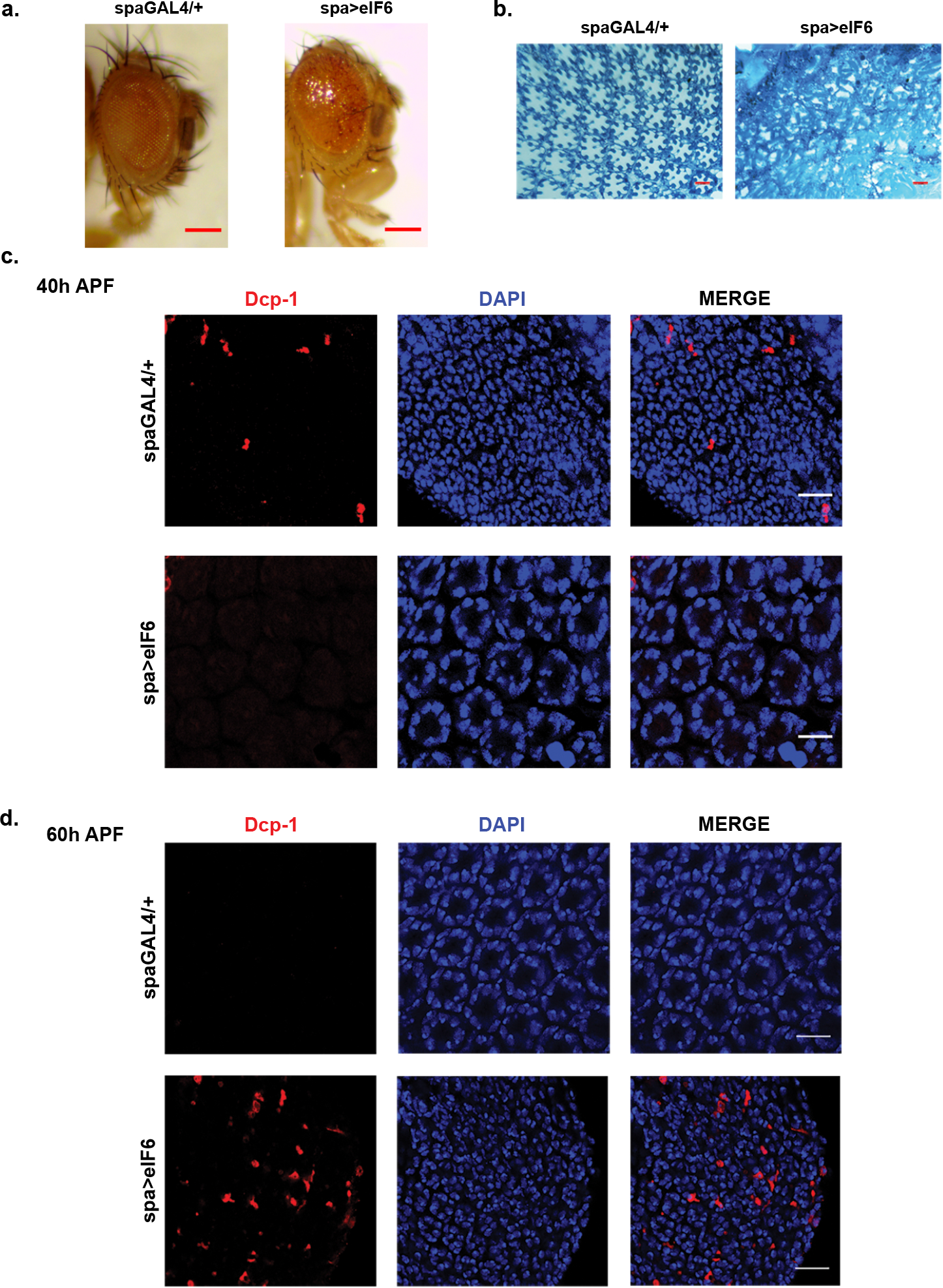
A specific increase of *eIF6* gene dosage in cone cells results in a rough eye phenotype. **(a-b)** Overexpression of eIF6 in cone cells results in rough eye phenotype. **(a)** Representative stereomicroscope images of *spaGAL4/+* and *spa>eIF6* eyes showing a rough eye phenotype. Scale bar 100 µm **(b)** Representative tangential semithin sections of *spaGAL4/+* and *spa>eIF6* adult eyes showing disruption of the structure upon eIF6 overexpression in cone cells. Scale bar 10 µm. **(c)** Mid-pupal stage (40h APF) retinae of *spaGAL4/+* and *spa>eIF6* genotypes stained for Dcp-1 confirm the block in apoptosis already demonstrated in *GMR>eIF6* retinae. **(d)** Late-pupal stage (60h APF) retinae of *spaGAL4/+* and *spa>eIF6* genotypes stained for Dcp-1 confirming the delayed and increased apoptosis already observed in *GMR>eIF6* retinae. (c-d) Scale bar 10 µm.

### eIF6 expression reshapes the transcriptome, increasing a ribosome signature and repressing ecdysone signaling

Next, we asked whether eIF6 was associated with a transcriptional rewiring that could account for the observed phenotypic effects. To this end, we performed a comprehensive gene expression analysis of *GMRGAL4/+* and *GMR>eIF6* genotypes at two distinct stages of eye development, larval eye imaginal discs and pupal retinae by RNA-Seq (Fig. 5). In *GMR>eIF6* samples at both developmental stages, we observed an upregulation of genes involved in ribosome biogenesis (Fig. 5a, Supplementary File 1). GSAA analysis revealed also an increase in mRNAs of genes involved in rRNA processing (Fig. 5c). Overall these data suggest that eIF6 is able to increase ribosomal gene expression.

**Figure 5.**
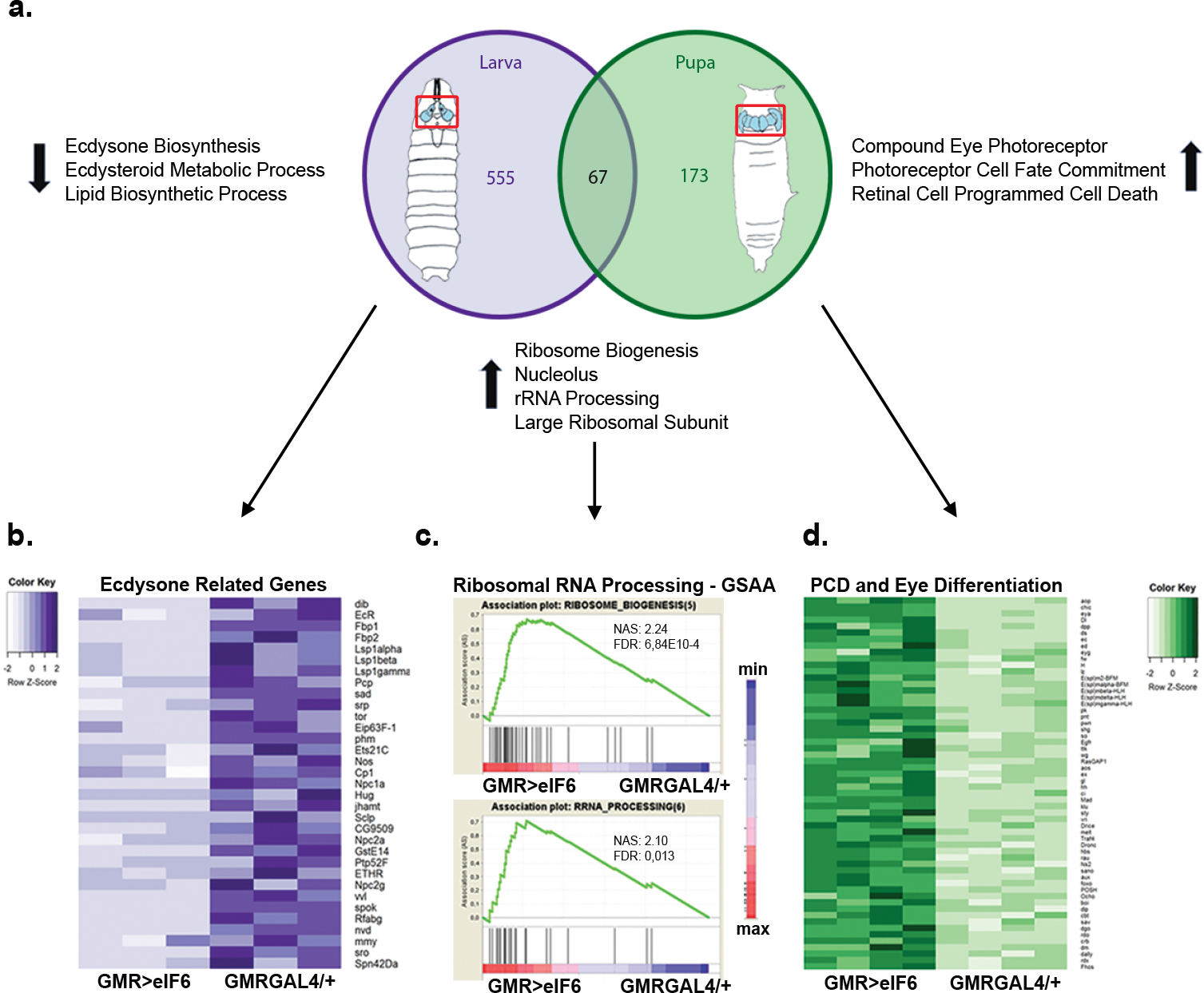
eIF6 induces a reshaping of transcription, resulting in rRNA processing alteration and in a gene signature specific for the eye. **(a)** Venn Diagram indicating genes differentially expressed in *GMR>eIF6* larval eye imaginal discs and *GMR>eIF6* retinae with respect to controls (*GMRGAL4/+)*. **(b)** The Ecdysone Biosynthetic Pathway is shut off when eIF6 is upregulated. Heat Map representing absolute gene expression levels in *GMR>eIF6* and *GMRGAL4/+* eye imaginal disc samples for the subset of gene sets involved in Ecdysone Biosynthesis by Gene Ontology analysis. **(c)** Gene Set Association Analysis (GSAA) indicates a significative upregulation of ribosomal machinery. Representative Enrichment Plots indicating a striking upregulation of genes involved in rRNA Processing and Ribosome Biogenesis in both *GMR>eIF6* eye imaginal discs and *GMR>eIF6* retinae with respect to their controls (*GMRGAL4/+)*. **(d)** mRNAs involved in Programmed Cell Death and in Eye Differentiation are upregulated in *GMR>eIF6* retinae. Heat Map representing absolute gene expression levels in *GMR>DeIF6* and *GMRGAL4/+* retinae samples for the subset of gene sets involved in Programmed Cell Death and Eye Differentiation by Gene Ontology Analysis.

Consistent with our phenotypic analysis of the eye, *GMR>eIF6* retinae displayed also variations in genes involved in eye development and in PCD (Fig. 5a, d and Supplementary File 1). Notably, mRNAs encoding specialized eye enzymes, such as those of pigment biosynthetic pathways, were downregulated in *GMR>eIF6* samples (Supplementary File 1), preceding the altered adult eye morphology.

Finally, coordinated changes induced by eIF6 in eye imaginal discs surprisingly clustered into the ecdysone pathway, with a striking downregulation of many enzymes involved in 20-HydroxyEcdysone (20-HE) biosynthesis (Fig. 5 a-b). For instance, expression of *phm*, *sad* and *nvd* (Supplementary Fig. 5a) were virtually absent in *GMR>eIF6* eye imaginal disc, while early (*rbp*) and late (*ptp52f*) responsive genes belonging to the hormone signaling cascade were downregulated (Supplementary File 1). In conclusion, our gene expression analysis of *GMR>eIF6* eye samples identifies a rewiring of transcription that is consistent with altered PCD, accompanied by upregulation of ribosomal genes and downregulation of the ecdysone biosynthetic pathway.

### Increasing *eIF6* gene dosage results in elevated translation

eIF6 binds free 60S *in vitro* and *in vivo* affecting translation ^7^. To assess whether increased transcription of genes related to ribosome biogenesis and rRNA processing observed in gene expression analysis experiments was accompanied by an effect in the translational machinery, we investigated changes in levels of free 60S subunits upon eIF6 overexpression. To this end, we performed the *in vitro* Ribosome Interaction Assay (iRIA) ^33^. We found that the expression of eIF6 in larval eye discs (*GMR>eIF6*) led to a 25% reduction in free 60S sites when compared to control (*GMRGAL4/+*) (Fig 6a). Next, we used a modified SUnSET assay ^34^, as a proxy of the translational rate. We measured translation in eye imaginal discs treated *ex vivo* with puromycin, which incorporates in nascent protein chains by ribosomes. Remarkably, *GMR>eIF6* eye discs incorporated almost twice the amount of puromycin, relative to controls (Fig. 6b-c). Taken together, high levels of eIF6 increase the free 60S pool *in vivo*, and increase puromycin incorporation, i.e. translation. We next determined whether the observed effects were specific of eye development or rather a more general outcome associated with increased *eIF6* gene dosage. Thus, we overexpressed eIF6 in a different epithelial organ, the wing imaginal disc, using the *bxMS1096GAL4* driver (*MS>eIF6*). Such manipulation led to complete disruption of the adult wing structure (Fig. 6d and Supplementary Fig. 6c). Moreover, we performed the SUnSET assay on wing imaginal discs, and, as in eye discs, we observed a two-fold increase in puromycin incorporation in *MS>eIF6* wing discs with respect to the controls (Fig. 6e and Supplementary Fig. 6a). Finally, eIF6 overexpression in wing discs led to the presence of apoptotic cells in the dorsal portion of the disc (Supplementary Fig. 6b), as previously evidenced in the developing retina. In conclusion, the increased gene dosage of *eIF6* leads to augmented translational activity that results in a specific gene expression program.

**Figure 6.**
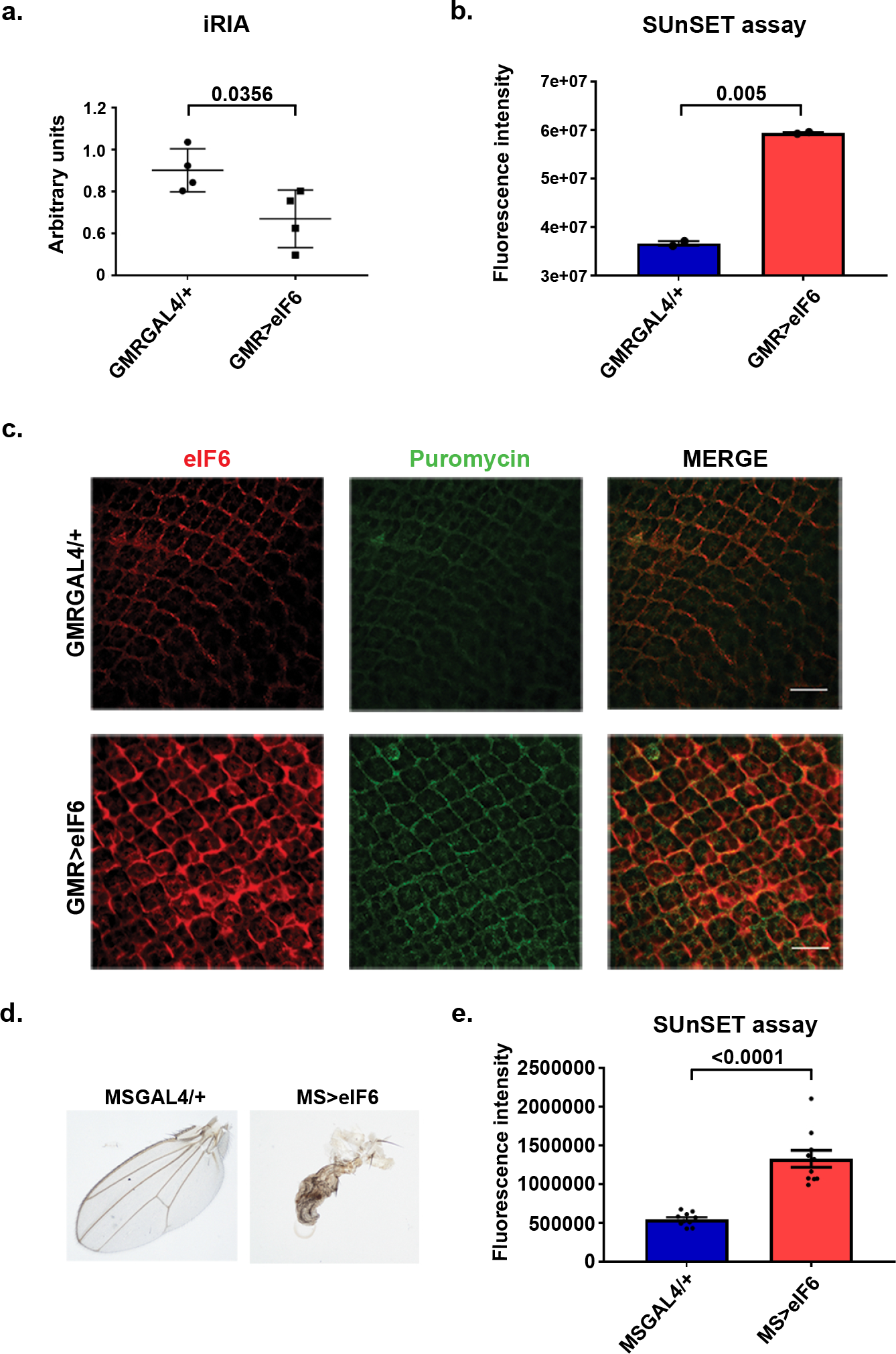
Increased eIF6 levels in the developing eye result in reduced free 60S and increased translation. **(a)** *In vitro* iRIA assays showing that *eIF6* increased dosage reduce the number of free 60S subunits. Values represent the mean ± SEM from two replicates. Assays were repeated three times. Student’s t-test was used to calculate p values. **(b)** *In vitro* SUnSET assays showing that *eIF6* increased gene is associated with increased puromycin incorporation. Barplots represent the mean ± SEM from three replicates. Assays were repeated three times. Student’s t-test was used to calculate p values. Quantification of SUnSET assay was performed with ImageJ software. **(c)** Representative SUnSET assay performed using immunofluorescence experiments, indicating a two-fold increase in general translation when eIF6 levels are increased in eye imaginal discs. Scale bar 10 µm. **(d)** Adult wings *MS>eIF6* have a completely aberrant phenotype. **(e)** *In vitro* SUnSET assays showing that *eIF6* increased gene is associated with 2-fold puromycin incorporation in wing discs. Barplots represent the mean ± SEM from three replicates. Assays were repeated three times. Student’s t-test was used to calculate p values. Quantification of SUnSET assay was performed with ImageJ software.

### 20-HE administration rescues adult eye defects induced by increased *eIF6* gene dosage

Transcriptome analysis revealed a coordinated shutdown of the 20-HE biosynthetic pathway raising the question whether 20-HE administration could at least partly rescue the defects driven by eIF6 increased levels, and a rough eye phenotype characterized by aberrant PCD. To determine the hierarchy of events that the increased *eIF6* gene dosage causes, we administrated the active form of the hormone 20-HE by feeding *GMR>eIF6* third instar larvae with 20-HE and we evaluated the effect on eye development, finding a partial rescue of the rough phenotype. Remarkably, *GMR>eIF6* larvae fed with 20-HE showed eyes that were 20% larger than untreated controls, although they remained smaller with respect to *GMRGAL4/+* (Fig. 7a). We also assessed the levels of apoptosis at 40h APF. Notably, immunofluorescence staining for Dcp-1 showed the presence of apoptotic cells in 40h APF *GMR>eIF6* retinae treated with 20-HE, while *GMR>eIF6* untreated retinae did not show any Dcp-1 positive cells (Fig. 7b). Taken toghether, these data suggest that the apoptotic defect and eye roughness caused by increased *eIF6* gene dosage are due to the inactivation of ecdysone signaling, that precedes a deregulation of PCD.

**Figure 7.**
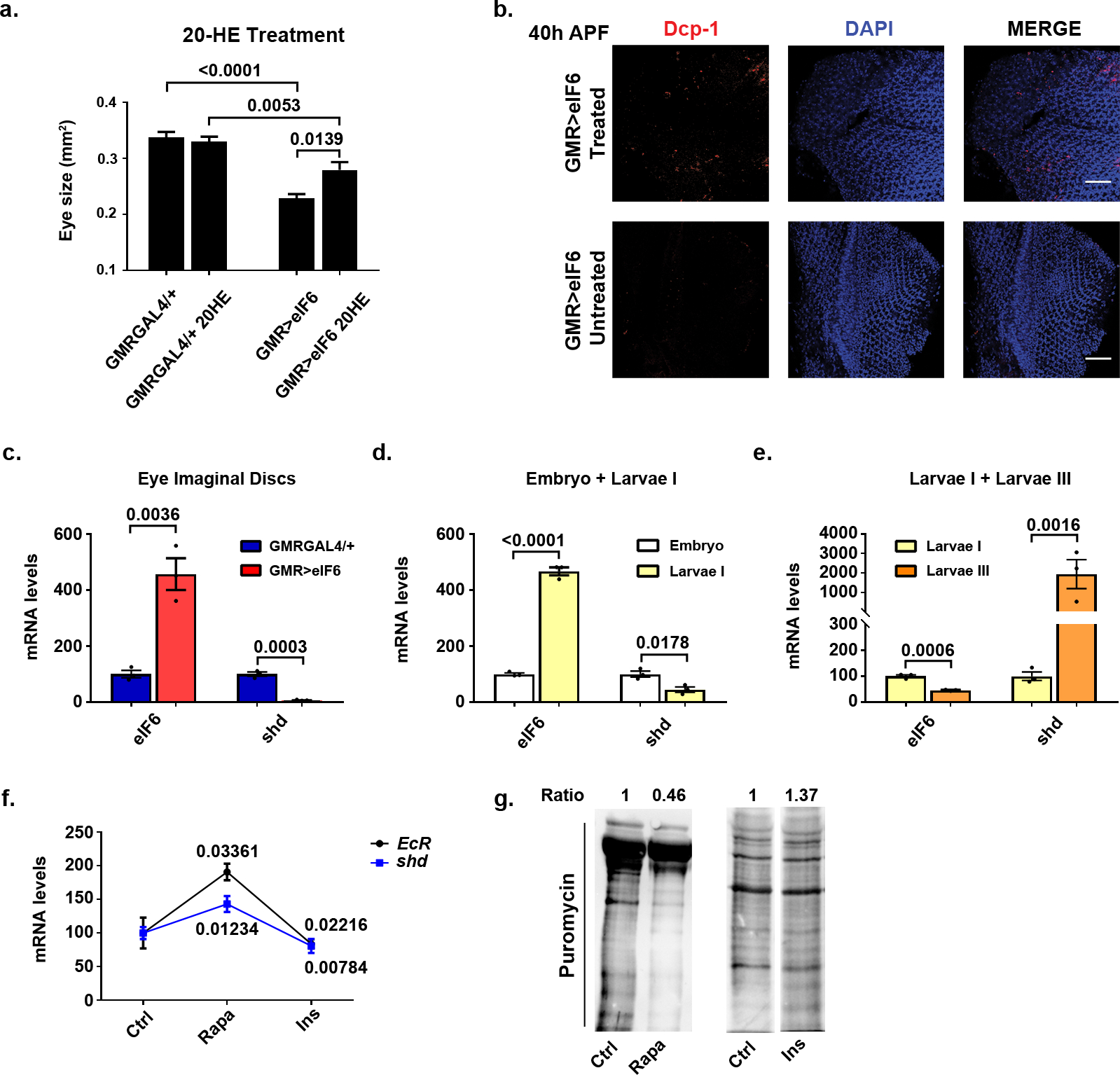
20-HE treatment rescues the rough eye phenotype due to high levels of eIF6, unveiling the role of translation in ecdysone biosynthesis regulation. **(a-b)** 20-HE treatment partially rescue the rough eye phenotype and the delay in apoptosis in 40h APF retinae **(a)** The barplot represents the average of n>8 independently collected samples with error bars indicating the SEM. P-values were calculated using an upaired two-tailed Student t-test. The graph shows the *GMR>eIF6* adult fly eye size with or without treatment with 20-HE. As indicated in the barplot, the fly eye size is partially rescued when the hormone is added to the fly food. **(b)** Immunofluorescence images showing that 20-HE treatment (240 µg/mL in standard fly food) rescues the apoptotic delay observed in *GMR>eIF6* 40h APF retinae. **(c-e)** Real-time PCR analyses of the indicated genes showing an inverse correlation between eIF6 and shd mRNA levels. The RNA level of each gene was calculated relative to RpL32 expression as a reference gene. The barplot represents the average of at least three independent biological replicates with error bars indicating the SEM. p-values were calculated using an upaired two-tailed Student t-test. **(c)** Real-time PCR analyses of the indicated genes in *GMRGAL4/+* and *GMR>eIF6* eye imaginal discs. Upon eIF6 overexpression, *GMR>eIF6* eye imaginal discs have less abundance of *shd* mRNA levels compared to *GMRGAL4/+* eye imaginal discs. **(d-e)** During development, *eIF6* and *shd* mRNA levels show an inverse correlation by comparing embryos with first instar larval RNA extracts **(d)** or by comparing first and thirs instar larval RNA extracts **(e)**. **(f-g)** The ecdysone biosynthetic pathway genes *shd* and *EcR* are modulated upon translation modulation in S2 cells. **(f)** Real time analysis evidences that upon inhibition of translation with rapamycin treatment (1 µM, 2 hours) the level of *shd* and *EcR* mRNA levels increase, contrary to the drop observed upon translation stimulation with insulin (1 µM, 12 hours). The RNA level of each gene was calculated relative to RpL32 expression as a reference gene. The barplot represents the average of at least three independent biological replicates with error bars indicating the SEM. p-values were calculated using an upaired two-tailed Student t-test. **(g).** Representative western blot showing the decreased or increased rate of protein synthesis upon rapamycin or insulin treatment respectively with SUnSET method ^34^

### eIF6 and translation antagonize ecdysone biosynthesis during development

Our findings indicate that increased eIF6 levels cause downregulation of mRNAs belonging to the ecdysone biosynthetic pathway, and the relative absence of its final product, the 20-HE. To understand the physiological relevance of this phenomenon, we measured mRNAs levels of *eIF6* and *shd* (as a proxy of the entire ecdysone biosynthetic pathway) which encodes for the last enzyme of the 20-HE biosynthesis, at different stages of development (Fig. 7). We first confirmed by Real-Time PCR the downregulation of *shd* in eye imaginal disc overexpressing eIF6 (Fig. 7c). We then investigated the levels of *eIF6* and *shd* during development in wild-type tissues (Fig. 7d-e). Interestingly, we found that eIF6 levels are regulated during development, and that *shd* levels drop when *eIF6* levels are high, both by comparing embryos and first instar larvae (Fig. 7d) or first and third instar larvae (Fig. 7e). Taken together, data suggest that *eIF6* gene dosage physiologically is inversely correlate with 20-HE production. To verify the inverse relationship between the translational rate and ecdysone production, we assessed levels of *shd* and *EcR* (as an index of the feed forward loop induced by 20-HE itself ^35^) mRNA levels in S2 cells after treatment with rapamycin or insulin to inhibit or stimulate translation respectively (Fig. 7f-g). After insulin treatment we observed the downregulation of *shd* and *EcR* mRNA levels (Fig. 7f). Conversely, after rapamycin treatment, which decreases mTORC1-regulated protein synthesis, we found an upregulation of the two analyzed genes (Fig. 7f). These data support a physiological model in which translation is a negative regulator of ecdysone metabolism.

## DISCUSSION

Eukaryotic Initiation Factor *eIF6* is an evolutionarily conserved gene encoding for a protein necessary for ribosome biogenesis and translation initiation ^8,9^. However, in mammals, eIF6 expression differs among tissues, with high levels in embryos and in cycling cells and almost undetectable levels in post-mitotic cells ^15^. Developmental studies in mice demonstrated that null alleles for this initiation factor are incompatible with life ^8^, whereas eIF6 haploinsufficiency is linked to an impairment in G1/S cell cycle progression ^8^. In unicellular models, eIF6 mutations rescue the quasi-lethal phenotype due to loss of ribosome biogenesis factors such as SBDS^12^. Taken together, these data highlight how eIF6 expression, despite of its ubiquitous function, is strictly regulated. Indeed, we found that doubling levels of eIF6 during development disrupts eye morphology, increases translation and changes profoundly gene expression. Overall, our data demonstrate that eIF6 is a translation factor able to drive a complex transcriptional reshaping.

Mechanistically, eIF6 binds to the 60S in the intersubunit space, interacting with rpL23 and to the sarcin-loop (SRL) of rpL24 ^36^, thus generating a steric hindrance that prevents the formation of a intersubunit bridge ^37^. *In vitro*, eIF6 can repress translation ^38^. In mice, however, high levels of eIF6 are required for both tumor progression ^18^, and insulin-controlled translation ^7,8^. In Drosophila, we found that an overexpression of eIF6 leads to a reduction of the free 60S pool in eye imaginal discs, consistent with eIF6 biochemical activity. Such reduction could imply lower general translation, due to less availability of 60S subunits, as in the case of Sbds mutants^39^. Conversely, 60S could be already engaged with 40S into active translating 80S, thus heightening general translation. We favor the latter hypothesis because by puromycin incorporation assay we see a two-fold increase in general translation, both in the developing eye and in the wing. Intriguingly, the transcriptome signature associated with high levels of eIF6 revealed also an increase in mRNAs encoding for rRNA processing factors, suggesting that ribosome biogenesis is positively affected by eIF6. In conclusion, we surmise that *in vivo* eIF6 may act as a powerful stimulator of ribosome synthesis and translation. The effects associated with increased translation driven by eIF6 are at least two, a change in the ecdysone pathway and a delay in apoptosis. We found a strong reduction of ecdysone biosynthesis and signaling pathway in eye imaginal disc driven by eIF6. 20-HE treatment leads to a partial rescue of the developmental defects driven by eIF6 increased activity. Thus, our data suggest that eIF6 is upstream of ecdysone regulation. It has been recently suggested how translation regulation and hormonal signaling are tightly interconnected in Drosophila ^40^ and, more generally, that translation is a controller of metabolism ^41^. Our experiments unveil an inverse correlation between translational capability and ecdysone production. Concerning apoptosis we showed that eIF6 expression leads to an early block in Programmed Cell Death, as previously demonstrated by others in *X. laevis* ^42^. The developmental defects driven by increased *eIF6* gene dosage are consistent with two scenarios: excess eIF6 could delay developmental PCD. Alternatively, PCD could be repressed at the correct developmental time and apoptotic elimination of defective cells overexpressing eIF6 could be triggered later independently of developmental signals. The fact that overexpression of eIF6 in wing discs, which are not subjected to developmental apoptosis, leads to cell death supports the latter hypothesis.

The developmental changes due to eIF6-driven translation are dramatic and include lethality, as well as disruption of development. In the past, similar effects were observed by the expression of another rate-limiting factor in translational initiation, eIF4E ^43^. It is unknown whether the developmental defects driven by eIF4E overexpression also included the arrest of ecdysone biosynthetic pathway, or an apoptotic block. However, in mammalian models eIF4E and eIF6 share the common property of being rate-limiting for tumor growth and translation in several contexts ^44-50^. In summary, our study demonstrates that overexpression of eIF6 in developing organs is sufficient to induce an increase in ribosome biogenesis and translation that correlates with complex transcriptional and metabolic changes leading to hormonal and apoptotic defects. It will be interesting to further dissect the relationship between epigenetic, metabolic, and transcriptional changes in the *Drosophila* model. Our model may also be useful for *in vivo* screenings of compounds that suppress the effect of eIF6 overexpression. Such approach could isolate useful therapeutics that might be relevant to the protumourigenic role of mammalian eIF6, and could identify novel genetic modulators of eIF6 function. In short, translation factors may drive complex and coordinated programs upstream of transcription.

## MATERIALS AND METHODS

### Genetics

Fly strains were maintained on standard cornmeal food at 18°C. Genetic crosses were performed at 25°C, with the exception of *GMRGAL/+* and *GMR>eIF6*, performed at 18°C. The following fly mutant stocks have been used: *GMRGAL4/CTG* was a gift from Manolis Fanto (King’s College, London); *UAS-eIF6* was a gift from William J Brook (Alberta Children’s Hospital, Calgary) ^51^. Lines obtained from the Bloomington Drosophila Stock Center (BDSC): *spaGAL4* (26656), *54CGAL4* (27328), *w1118*, *bxMS1096GAL4* (8860).

### Mosaic analysis

The *eIF6^k13214^* mutant clones were created by *Flippase* (*FLP*) mediated mitotic recombination ^31^. The *DeIF6^k13214^ (P(w[+mC]=lacW) eIF6[k13214] ytr[k13214]) P* element allele was recombined onto the right arm of chromosome two with the homologous recombination site (FRT) at 42D using standard selection techniques. Briefly, to create the FRT *y^+^ pwn, DeIF6^k13214^* chromosomes, *eIF6^k13214^* was recombined onto the FRT chromosome originating from the *y; P42D pwn[1] P{y+}44B/CyO* parental stock. The *yellow^+^ pwn DeIF6^k13214^G418* resistant flies were selected to create stocks for clonal analysis. Similarly, stocks used for generating unmarked *eIF6^k13214^* clones were created by recombining *eIF6^k13214^* with the 42D FRT chromosome using the *w[1118]; P42D P{Ubi-GFP}2R/CyO* parental line. Targeted mitotic wing clones were generated by crossing flies with *UAS-FLP*, the appropriate GAL4 driver and the suitable *42D* FRT second chromosome with the *42D* FRT *eIF6^k13214^*. The *hs* induced *eIF6^k13214^* mitotic clones were created by following standard techniques. Briefly, 24- and 48-hours larvae with the appropriate genotypes were heat shocked for 1 hour at 37°C followed by incubation at 25°C.

### S2 cell culture

The *Drosophila* S2 cells were grown in Schneider medium (Lonza, Basel, Switzerland, #04-351Q) supplemented with 10% Fetal Bovine Serum (FBS – #ECS0180L, Euroclone, Pero, Italy) and 5 mL of PSG 1X (100X composition: 10000 U/mL Penicillin, 10 mg/mL Streptomycin and 200 mM L-Glutamine in citrate buffer, (#G1146, Sigma, St. Louis, MO), and maintained as a semi-adherent monolayer at standard culture conditions at 25 °C without CO_2_. For protein synthesis measurement, S2 cells were treated at 65-70% confluence with 1 μM rapamycin (#R8781, Sigma) for 2 hours or 1 μM insulin (#I0516, Sigma) for 12 hours, both at 25 °C. For SUnSET assay, medium was removed and replaced with fresh medium supplemented with 5 μg/mL puromycin (#A1113803, Thermofisher Scientific, Waltham, MA, USA) for 3 hours, and treated according to ^34^.

### RNA isolation and RNA sequencing

Total RNA was extracted with mirVana^TM^ isolation kit according to the manufacturer protocols (#AM 1560, ThermoFisher) from 10 eye imaginal discs (larval stage) or 10 retinae (pupal stage). RNA quality was controlled with BioAnalyzer (Agilent, Santa Clara, CA, USA). Libraries for Illumina sequencing were constructed from 100 ng of total RNA with the Illumina TruSeq RNA Sample Preparation Kit v2 (Set A) (Illumina, San Diego, CA, USA). The generated libraries were loaded on to the cBot (Illumina) for clustering on a HiSeq Flow Cell v3. The flow cell was then sequenced using a HiScanSQ (Illumina). A paired-end (2×101) run was performed using the SBS Kit v3 (Illumina). Sequence deepness was at 35 million reads. For quantitative PCR, the same amount of RNA was retrotranscribed according to SuperScript^TM^ III First-Strand Synthesis SuperMix manufacturer protocol (#18080400, LifeTechnologies, Carlsbad, CA, USA). For RNA-Seq validation, Taqman probes specific for *eIF6* (Dm01844498_g1) and *rpl32* (Dm02151827_g1) were used, together with standard primers (*rpl32* Fwd CGGATCGATATGCTAAGCTGT, Rev CGACGCACTCYCYYGTCG; *shd* Fwd CGGGCTACTCGCTTAATGCAG, Rev AGCAGCACCACCTCCATTTC). Target mRNA quantification was performed by using ΔCt-method with *rpl32* RNA as an internal standard, performed on a StepOne Plus System (Applied Biosystems, Foster City, CA, USA).

### Bioinformatic Analysis

#### Read pre-processing and mapping

Three biological replicates were analyzed for *GMRGAL4/+* and *GMR>eIF6* larval eye imaginal discs and four biological replicates were analyzed for *GMRGAL4/+* and *GMR>eIF6* pupal retinae, for a total of 14 samples. Raw reads were checked for quality by FastQC software (version 0.11.2, S., A. FastQC: a quality control tool for high-throughput sequence data. 2010; Available from: http://www.bioinformatics.babraham.ac.uk/projects/fastqc), and filtered to remove low quality calls by Trimmomatic (version 0.32) ^52^ using default parameters and specifying a minimum length of 50. Processed reads were then aligned to *Drosophila melanogaster* genome assembly GRCm38 (Ensembl version 79) with STAR software (version 2.4.1c) ^53^.

#### Gene expression quantification and differential expression analysis

HTSeq-count algorithm (version 0.6.1, option -s = no, gene annotation release 79 from Ensembl) ^54^ was employed to produce gene counts for each sample. To estimate differential expression, the matrix of gene counts produced by HTSeq was analyzed by DESeq2 (version DESeq2_1.12.4) ^55^. The differential expression analysis by the DeSeq2 algorithm was performed on the entire dataset composed by both larvae and pupae samples. The two following comparisons were analyzed: *GMR>eIF6 versus GMRGAL4/+* larval eye imaginal discs (6 samples overall) and *GMR>eIF6 versus GMRGAL4/+* pupal retinae (8 samples in total). Reads counts were normalized by calculating a size factor, as implemented in DESeq2. Independent filtering procedure was then applied, setting the threshold to the 62 percentile; 10886 genes were therefore tested for differential expression. Significantly modulated genes in *GMR>eIF6* genotype were selected by considering a false discovery rate lower than 5%. Regularized logarithmic (rlog) transformed values were used for heat map representation of gene expression profiles. Analyses were performed in R version 3.3.1 (2016-06-21, Computing, T.R.F.f.S. R: A Language and Environment for Statistical Computing. Available from: http://www.Rproject.org/).

#### Functional analysis by topGO

The Gene Ontology enrichment analysis was performed using topGO R Bioconductor package (version topGO_2.24.0). The option *nodesize = 5* is used to prune the GO hierarchy from the terms which have less than 5 annotated genes and the *annFUN.db* function is used to extract the gene-to-GO mappings from the genome-wide annotation library *org.Dm.eg.db* for *D. melanogaster*. The statistical enrichment of GO was tested using the Fisher’s exact test. Both the “classic” and “elim” algorithms were used.

#### Gene set association analysis

Gene set association analysis for larvae and pupae samples was performed by GSAA software (version 2.0) ^56^. Raw reads for 10886 genes identified by Entrez Gene ID were analyzed by GSAASeqSP, using gene set C5 (*Drosophila* version retrieved from http://www.go2msig.org/cgi-bin/prebuilt.cgi?taxid=7227) and specifying as permutation type ‘gene set’ and as gene set size filtering min 15 and max 800.

### Western blotting and antibodies

Larval imaginal discs, pupal retinae and adult heads were dissected in cold Phosphate Buffer Saline (Na_2_HPO_4_ 10 mM, KH_2_PO_4_ 1.8 mM, NaCl 137 mM, KCl 2.7 mM, pH 7.4) (PBS) and then homogenized in lysis buffer (HEPES 20 mM, KCl 100 mM, Glycerol 5%, EDTA pH 8.0 10 mM, Triton-X 0.1%, DTT 1mM) freshly supplemented with Protease Inhibitors (Sigma, St. Louis, MO, USA, #P8340). Protein concentration was determined by BCA analysis (Pierce, Rockford, IL, USA, #23227). Equal amounts of proteins were loaded and separated on a 10% SDS-PAGE, then transferred to a PVDF membrane. Membranes were blocked in 10% Bovine Serum Albumin (BSA) in PBS-Tween (0.01%) for 30 minutes at 37°C. The following primary antibodies were used: rabbit anti-eIF6 (1:500, this study), rabbit anti-β-actin (1:4000, CST, Danvers, MA, USA, #4967), mouse anti-Puromycin (1:500, Merck Millipore, #MABE343). To produce the anti-eIF6 antibody used in this study, a rabbit polyclonal antiserum against two epitopes on COOH-terminal peptide of eIF6 (NH2-CLSFVGMNTTATEI-COOH eIF6 203-215 aa; NH2-CATVTTKLRAALIEDMS-COOH eIF6 230-245 aa) was prepared by PrimmBiotech (Milan, Italy, Ab code: 201212-00003 GHA/12), purified in a CNBr-Sepharose column and tested for its specificity against a mix of synthetic peptides with ELISA test. The following secondary antibodies were used: donkey anti-mouse IgG HRP (1:5000, GE Healthcare, Little Chalfont, UK, Amersham #NA931) and donkey anti-rabbit IgG HRP (1:5000, GE Healthcare, Amersham #NA934).

### SUnSET Assay

Larval imaginal eye and wing discs were dissected in complete Schneider medium (Lonza, Basel, Switzerland) and treated *ex vivo* with puromycin (50 µg/mL) for 30 minutes at room temperature, then fixed in 3% paraformaldehyde (PFA) for 1 hour at room temperature. Immunofluorescences were then performed as described below, using a mouse anti-puromycin (1:500, Merck Millipore, Billerica, MA, USA, #MABE343) as a primary antibody. Discs were then examined by confocal microscope (Leica SP5, Leica, Wetzlar, Germany) and fluorescence intensity was measured with ImageJ software.

### Cells count

*GMRGAL4/+* and *GMR>eIF6* pupal retinae at 40h APF were dissected, fixed, and stained with anti-Armadillo to count cells, as previously described ^57^. Cells contained within a hexagonal array (an imaginary hexagon that connects the centers of the surrounding six ommatidia) were counted; for different genotypes, the number of cells per hexagon was calculated by counting cells, compared with corresponding control. Cells at the boundaries between neighboring ommatidia count half. At least 3 hexagons (equivalent to 9 full ommatidia) were counted for each genotype, and phenotypes were analysed. Standard Deviation (SD) and unpaired two-tailed Student t-test were used as statistical analysis.

### Immunofluorescences, antibodies and TUNEL Assay

Larval imaginal discs and pupal retinae were dissected in cold PBS and fixed in 3% paraformaldehyde (PFA) for 1 hour at room temperature, then washed twice with PBS and blocked in PBTB (PBS, Triton 0.3%, 5% Normal Goat Serum and 2% Bovine Serum Albumin) for 3 hours at room temperature. Primary antibodies were diluted in PBTB solution and incubated O/N at 4°C. After three washes with PBS, tissues were incubated O/N at 4°C with secondary antibodies and DAPI (1:1000, Molecular Probes, Eugene, OR, USA, #D3571) in PBS. After three washes with PBS, eye imaginal discs and retinae were mounted on slides with ProLong Gold (LifeTechnologies, Carlsbad, CA, USA, #P36930). The following primary antibodies were used: rabbit anti-eIF6 (1:50, this study), rat anti-ELAV (1:100, Developmental Study Hybridoma Bank DSHB, Iowa City, IA, USA, #7E8A10), mouse anti-CUT (1:100, DSHB, #2B10), mouse anti-Armadillo (1:100, DSHB, #N27A), mouse anti-Chaoptin (1:100, DSHB, #24B10), rabbit anti-Dcp-1 (1:50, CST, #9578), mouse anti-Puromycin (1:500, Merck Millipore, #MABE343). The following secondary antibodies were used: donkey anti-rat, donkey anti-mouse, donkey anti-rabbit (1:500 Alexa Fluor^®^ secondary antibodies, Molecular Probes). Dead cells were detected using the In Situ Cell Death Detection Kit TMR Red (Roche, Basel, Switzerland, #12156792910) as manufacturer protocol, with some optimization. Briefly, retinae of the selected developmental stage were dissected in cold PBS and fixed with PFA 3% for 1 hour at room temperature. After three washes in PBS, retinae were permeabilized with Sodium Citrate 0.1%-Triton-X 0.1% for 2 minutes at 4°C and then incubated overnight at 37°C with the enzyme mix. Retinae were then rinsed three times with PBS, incubated with DAPI to stain nuclei and mounted on slides. Discs and retinae were examined by confocal microscopy (Leica SP5) and analysed with Volocity 6.3 software (Perkin Elmer, Waltham, MA, USA).

### Semithin sections

Semithin sections were prepared as described in ^58^. Adult eyes were fixed in 0.1 M Sodium Phosphate Buffer, 2% glutaraldehyde, on ice for 30 min, then incubated with 2% OsO_4_ in 0.1 M Sodium Phosphate Buffer for 2 hours on ice, dehydrated in ethanol (30%, 50%, 70%, 90%, and 100%) and twice in propylene oxide. Dehydrated eyes were then incubated O/N in 1:1 mix of propylene oxide and epoxy resin (Sigma, Durcupan™ ACM). Finally, eyes were embedded in pure epoxy resin and baked O/N at 70°C. The embedded eyes were cut on a Leica UltraCut UC6 microtome using a glass knife and images were acquired with a 100X oil lens, Nikon Upright XP61 microscope (Nikon, Tokyo, Japan).

### Ecdysone treatment

For ecdysone treatment, 20-HydroxyEcdysone (20HE) (Sigma, #H5142) was dissolved in 100% ethanol to a final concentration of 5 mg/mL; third instar larvae from different genotypes (*GMRGAL4/+* and *GMR>eIF6*) were collected and placed in individual vials on fresh standard cornmeal food supplemented with 240 µg/mL 20-HE. Eye phenotype was analyzed in adult flies, and images were captured with a TOUPCAM™ Digital camera. Eye images were analyzed with ImageJ software.

### *In vitro* Ribosome Interaction Assay (iRIA)

iRIA assay was performed as described in ^33^. Briefly, 96-well plates were coated with a cellular extract diluted in 50 µL of PBS, 0.01% Tween-20, O/N at 4°C in humid chamber. Coating solution was removed and aspecific sites were blocked with 10% BSA, dissolved in PBS, 0.01% Tween-20 for 30 minutes at 37 °C. Plates were washed with 100 μL/well with PBS-Tween. 0.5 μg of recombinant biotinylated eIF6 were resuspended in a reaction mix: 2.5 mM MgCl_2_, 2% DMSO and PBS-0.01% Tween, to reach 50 µL of final volume/well, added to the well and incubated with coated ribosomes for 1 hour at room temperature. To remove unbound proteins, each well was washed 3 times with PBS, 0.01% Tween-20. HRP-conjugated streptavidin was diluted 1:7000 in PBS, 0.01% Tween-20 and incubated in the well, 30 minutes at room temperature, in a final volume of 50 µL. Excess of streptavidin was removed through three washes with PBS-Tween. OPD (o-phenylenediamine dihydrochloride) was used according to the manufacturer’s protocol (Sigma-Aldrich) as a soluble substrate for the detection of streptavidin peroxidase activity. The signal was detected after the incubation, plates were read at 450 nm on a multiwell plate reader (Microplate model 680, Bio-Rad, Hercules, CA, USA).

### Data availability

Data generated by our RNASeq experiment have been deposited in ArrayExpress. Accession Number ID will be provided upon acceptance for publication.

## SUPPLEMENTARY DATA

For review, Supplementary Figures and Supplementary Files are available.

## ACKNOWLEDGEMENTS

We thank William Brook (Alberta Children’s Hospital, Calgary) for UASDeIF6 stocks and Manolis Fanto (King’s College, London) for stocks and suggestions. We thank Valeria Berno for imaging help and Vera Giulia Volpi for semithin sections preparation. This work was supported by ERC TRANSLATE 338999 and FONDAZIONE CARIPLO to SB. P.C. is supported by Fondazione Umberto Veronesi.

## AUTHOR CONTRIBUTIONS

A.R., G.G., E.P., K.S., S.R., M.M., P.C. performed experiments; A.R., G.G., P.C. prepared the manuscript and figures; T.V., S.B. and P.C. reviewed and edited the manuscript; R.A. and C.C. performed bioinformatic analysis; A.R., S.B. and P.C. conceived the project and experiments. All authors contributed, read, and approved the manuscript. The authors declare no competing interests.

